# Trimeric SARS-CoV-2 Spike proteins produced from CHO-cells in bioreactors are high quality antigens

**DOI:** 10.1101/2020.11.15.382044

**Authors:** Paco Pino, Joeri Kint, Divor Kiseljak, Valentina Agnolon, Giampietro Corradin, Andrey V. Kajava, Paolo Rovero, Ronald Dijkman, Gerco den Hartog, Jason S. McLellan, Patrick O. Byrne, Maria J Wurm, Florian M Wurm

## Abstract

The Spike protein of SARS-CoV-2 is essential for virus entry into human cells. In fact, most neutralizing antibodies against SARS-CoV-2 are directed against the Spike, making it the antigen of choice for use in vaccines and diagnostic tests. In the current pandemic context, global demand for Spike proteins has rapidly increased and could exceed hundreds of grams to kilograms annually. Coronavirus Spikes are large, heavily glycosylated, homotrimeric complexes, with inherent instability. Their poor manufacturability now threatens availability of these proteins for vaccines and diagnostic tests. Here, we outline a scalable, GMP-compliant, chemically defined process for production of a cell secreted, stabilized form of the trimeric Spike protein. The process is chemically defined and based on clonal, suspension-CHO cell populations and on protein purification via a two-step, scalable downstream process. The trimeric conformation was confirmed using electron microscopy and HPLC analysis. Binding to susceptible cells was shown using a virus-inhibition assay. The diagnostic sensitivity and specificity for detection of serum SARS-CoV-2 specific IgG1 was investigated and found to exceed that of Spike fragments (S1 and RBD). The process described here will enable production of sufficient high-quality trimeric Spike protein to meet the global demand for SARS-CoV-2 vaccines and diagnostic tests.

## 1. Introduction

Severe acute respiratory syndrome coronavirus 2 (SARS-CoV-2) is the virus responsible for the 2019 coronavirus disease (COVID-19) pandemic [1, 2] which presents an unprecedented challenge to societies globally. Effective vaccines and sensitive diagnostic tools for COVID-19 are urgently needed, and systems to produce and deliver these tools in sufficient quantities are required. It has become clear that contemporary protein production technologies for Spike proteins are insufficient to meet the unprecedented global demand for critical ingredients. As the SARS-CoV-2 trimeric Spike complex is a major target of the immune system, it could be a useful ingredient for vaccine and diagnostic applications. Stabilized trimeric Spike proteins have been selected as antigen of choice for RNA and virus-vector based vaccine candidates that are currently under development (Moderna, Novavax, Pfizer, J&J etc.). The efficacy of these vaccines is hoped or predicted to be between 50 and 70%. To boost the immune response, it is likely that follow-up vaccinations will be required. An adjuvated, highly purified subunit vaccine based on the stabilized trimeric Spike protein could be ideal for this purpose. Subunit vaccines can be produced in a cost-effective way and can be transported and stored lyophilized at ambient temperature. Also, whether any vaccination approach has elicited a sufficiently protective immune response against SARS-CoV-2 and the perpetuity of the immune response needs to be verified. To address this, the quantification of antibody levels against SARS-CoV-2 in serum is the most practical approach. For this reason, there will be a high demand for diagnostic tests for quantification of SARS-CoV-2-specific antibodies. The antigen which provides the best sensitivity and specificity for detection of SARS-CoV-2 antibodies is the trimeric form of the Spike protein [3].

The viral Spike (S) protein complex is a surface-exposed homo-trimeric structure that mediates entry into host cells. The Spike engages the cellular host receptor ACE2 and mediates virus-host membrane fusion. It is critical in the viral replication cycle and thus the Spike complex is considered the primary target of neutralizing antibodies [4–7]. The ideal diagnostic test for SARS-CoV-2 antibodies would detect all antibodies directed against the trimeric S protein complex. Production of such diagnostic tests implies the production of large quantities of highly purified S protein similar to its natural prefusion conformation [8]. The coronavirus Spike protein is a heavily glycosylated complex, with inherent structural flexibility and instability. In addition, the S protein is processed by Furin and the membrane-bound protease TMPRSS2 [6, 8]. Poor yield and pre-fusion instability of the S protein have hampered its use for development of vaccines and diagnostic tests. We explored the manufacturability of trimeric S protein in a secreted, soluble form using widely applied Chinese Hamster Ovary (CHO) cells. We developed a process that produces high quality S trimer protein for diagnostic tests as well as for vaccine applications. We used the fully characterized and CMC-compliant CHOExpress^™^ cell host (ExcellGene SA), single use equipment, chemically defined media and additives. Also, regulatory requirements from DNA construction to production in bioreactors were strictly followed. The clonally derived cell line and the scalable production process outlined here, will allow manufacture of trimeric S protein in grams and even kilograms, should the demand rise to such a level.

## 2. Material and Methods

### 2.1. Design of SARS CoV-2 Spike proteins

SARS-CoV-2 Spike protein encoding DNA were constructed and codon-optimized for CHO cells according to manufacturer recommendations (https://www.excellgene.com). A total of 11 DNA constructions were designed (Figure 1) and synthetized by ATUM (Menlo Park, USA). An IgG leader sequence was added to mediate efficient signal peptide cleavage. Histidine (8mer) encoding DNA was added for carboxyterminal expression. Where mentioned, the furin cleavage site RRAR is mutated to be non-functional. The transmembrane domain and the C terminal intracellular tail were removed and replaced by a T4 foldon sequence [9] in trimer designs. For the RBD-fragment (aa319-541) IgG leader and His-tag sequences were added. After appearance in infected patients of the D614G mutation in Spike proteins of the virus, this mutation was also incorporated into the chosen best construct.

**Figure 1:**
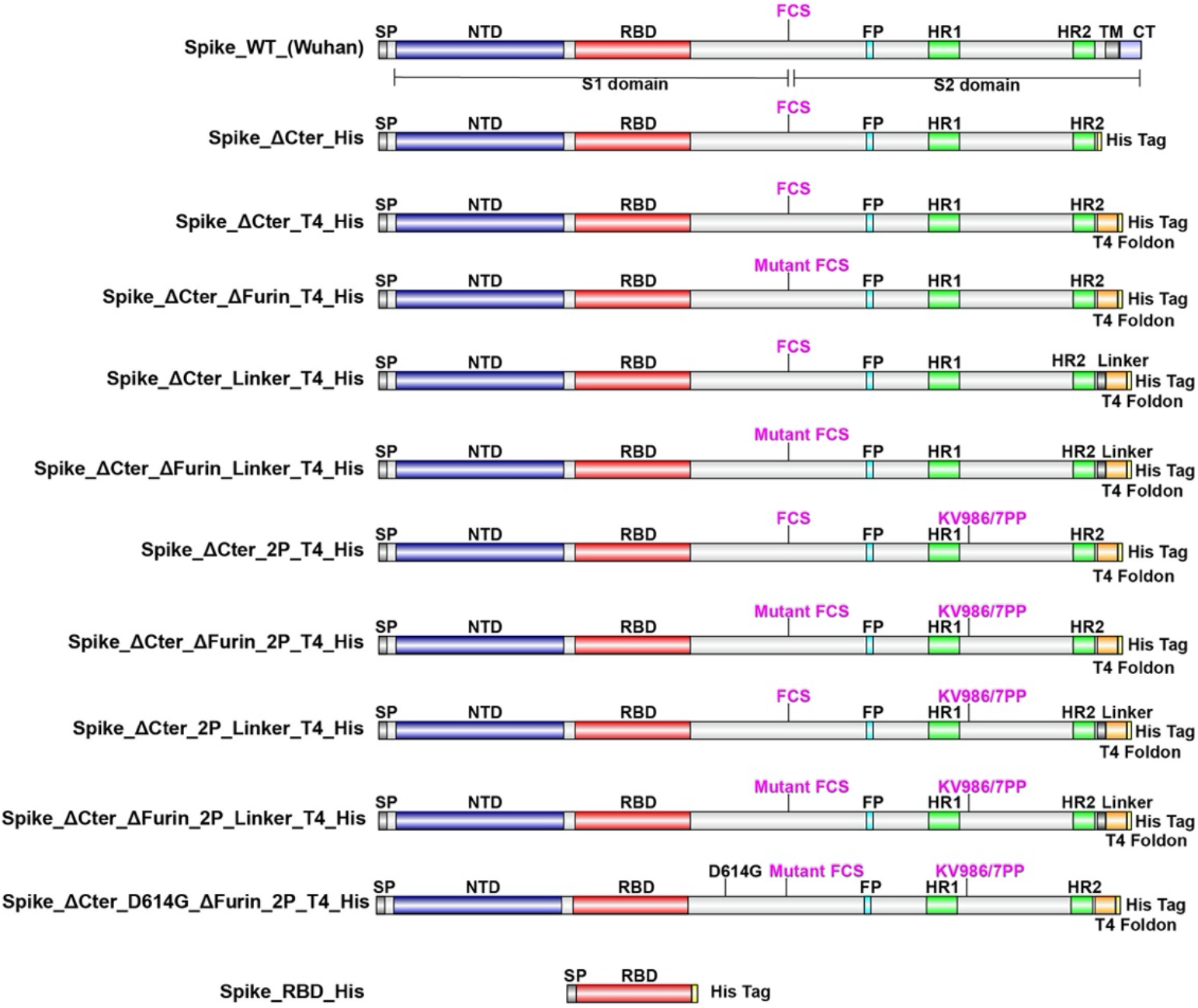
Schematic representation of SARS-CoV-2 Spike protein designs. SP: Signal Peptide, NTD: N Terminal Domain, RBD: Receptor Binding Domain, FCS: Furin Cleavage Site, FP: Fusion Peptide, HR1: Heptad Repeat 1, HR2: Heptad Repeat 2, TM: Transmembrane Domain, CT: C terminal Tail.

### 2.2 Expression and purification of SARS-CoV-2 proteins

Manufacturer’s protocols were followed for transfection and culturing of CHOExpress^™^ cells (ExcellGene SA, Monthey). An optimized transfection reagent mix (CHO4Tx^®^, ExcellGene SA) was used to transfect cells in animal-component free media with the vector pXLG-6 (ExcellGene SA) containing SARS-CoV-2 Spike DNA sequences, driven by promoter/enhancer sequences and associated elements. The vector includes an expression cassette for a puromycin resistance maker. Supernatants of high-density transient transfections under suspension culture were harvested after 10 days, mimicking a fed-batch process with ExcellGene’s animal component-free and protein-free CHO4Tx^®^ PM (production medium) in 50 mL TubeSpin^®^ bioreactor 50 tubes (TPP, Trasadingen, Switzerland). Cultures were shaken in a humidified ISF1-X/ISF3-X shaker (Kuhner, Birsfelden, Switzerland). Cultures were shifted from 37°C to 31°C during the production phase. Supernatants

Recombinant CHO cell lines: After transfections, suspension cells were stringently selected with puromycin. Resulting stable pools of recombinant CHO cells with satisfying yields were kept. Clonal cell lines were obtained by image-assisted cell distribution into single wells of 96 well plates (f-sight, Cytena GmbH, Freiburg). Expanded cell populations with high level expression for expected trimers and RDB monomers were frozen in mini-banks. Both, a recombinant pool cell line for the RBD Spike fragment and the lead clonal cell line for the Spike trimer were used for scale-up in an optimized fed-batch process at the 0.2 L, 10 L and 50 L bioreactor scale of operation.

The basic production medium employed EX-CELL^®^ Advanced^™^ CHO Fed-batch medium (Merck-Sigma). Bioreactors were seeded at 5×10^5^ cells/mL at 37°C. On day 4 the cell culture was shifted to 31° C, and animal component free 7a and 7b feeds (HyClone Cell Boost, Cytiva) were used according to an ExcellGene optimized procedure. Production culture fluids were harvested, clarified and subjected to purification by affinity chromatography after pumping them through layered (5 μm, 0.6μm, 0.2 μm) harvest filters (Cytiva, Ultrapure) to remove cells. Loading onto, washing of and elution from a Ni-Sepharose column (Cytiva) were optimized, following the resin producers’ suggestions. The eluted product stream was loaded on a Size-Exclusion column (SEC, Superdex 200 pg, Cytiva) for further purification, following a tangential flow filtration with a cut-off of 100’000 Dalton for the S trimer product. This step was omitted for the RBD fragment.

### 2.3. Electronmicroscopy analysis of Trimeric Spikes

Frozen aliquots of SARS-CoV-2 Spike trimer (Wuhan) and D614G mutant S trimer were thawed and purified by size exclusion chromatography (Superose 6 Increase 10/300, GE Healthcare) using 2 mM Tris pH 8.0, 200 mM NaCl and 0.02% (w/v) NaN_3_. Elution fractions containing S trimer were collected and diluted to 0.05 mg/mL. Samples were deposited onto plasma-cleaned, carbon-coated copper grids (CF400 mesh, Electron Microscopy Sciences), and stained in 2% (w/v) uranyl acetate at pH 7.0. Grids were imaged at a resolution of 92.000X in a Talos F200C transmission electron microscope equipped with a Ceta 16M detector (ThermoFisher Scientific). The pixel size was 1.63 Å. Contrast transfer function estimation and particle picking were performed in cisTEM [16]. Extracted particles were exported to cryoSPARC-v2 (Structura Biotechnology Inc.) for 2D classification, *ab initio* 3D reconstruction and homogeneous refinement. Three-fold symmetry (C3) was imposed during the final round of refinement.

### 2.4. Inhibition of SARS-CoV-2 infection by SARS-CoV-2-proteins

Briefly, SARS-CoV-2 virus (MOI of 0.01) was mixed with cell culture medium containing two-fold serial dilutions of RBD or trimeric Spike, ranging in concentration from 240 μM to 0.244 μM. This mixture was inoculated onto Vero E6 cells were inoculated with SARS-CoV-2 and at 48 hours post infection, SARS-CoV-2 infection was visualized using virus nucleocapsid antigen specific staining (red) and by a cell nucleus specific staining (blue), as described in [10]. Quantification of fluorescence was done as described [11].

### 2.5. Detection of SARS-CoV-2 antibodies using SARS-CoV-2-proteins

A bead-based serological assay [11] was used. Patient sera were collected and reacted with commercially available S-RBD and monomeric S1 domain protein, as well as S trimer and RBD. Briefly, 11ug of S-RBD (Sino Biologics, 40592-V08H), RBD (ExcellGene), monomeric Spike S1 (Sino Biologics, 40591-V08H) or S Trimer (ExcellGene), were loaded on 100 *μ*L of Microplex fluorescent beads and reacted with IgG containing sera. Relative optical readings were taken for each protein specific assay using the control sera and the COVID 19 sera. The relative reading for individual samples ranged from 0.01 AU/mL to about 1000 AU/mL. The sensitivity and specificity of the assay for each of the tested proteins was plotted in a Receiver Operating Characteristic (ROC) plot.

## 3. Results and discussion

### 3.1. Design and selection of CHO manufacturable SARS CoV-2 Trimeric Spikes

In order to define the optimal construct design to produce a soluble trimeric Spike protein we evaluated the expression with various S protein-variants (Figure 1) by CHO transient expression. The transmembrane domain and the C terminal intracellular tail were removed and replaced by a T4 foldon DNA sequence [9] with or without (GGGS)_n_ linker and a 8xHis tag encoding DNA. The furin cleavage site RRAR is mutated to be non-functional. WT Spike and amino-acid mutations K986P/V987P (“2P”) for locking the protein into a prefusion conformation [12] were also used. In addition, a receptor binding domain truncation of the S trimer was used (RBD-His). S proteins variants with a scrambled furin cleavage site were found better expressed. Mutation of the furin cleavage site seemed to increase trimer assembly and/or stability, suggesting that trimer assembly via T4 foldon occurs in the Golgi (Trans-Golgi network where furin is active). The same construct modifications were also applied for the D614G mutation in the Spike protein which has recently replaced to a large extent the Wuhan version.

Based on the relative expression levels in transient expressions with CHO cells combined with the efficiency of trimerization (Table 1), the Spike_ΔCter_ΔFurin_2P_T4_His design was selected for further evaluation. This design will be referred to in the following as S trimer.

**Table 1.** Overview of the relative expression levels of the different SARS-CoV-2 Spike protein designs evaluated under fed batch conditions, as well as their ability to form trimers in culture supernatants. ΔCter in the table refers to a construct that has the transmembrane and intracellular (TM, CT) section of the protein deleted.

| Construct | CHO Expression | Trimer Formation |
| --- | --- | --- |
| Spike_ΔCter_His | -- | n.a. |
| Spike_ΔCter_T4_His | - | + |
| Spike_ΔCter_ΔFurin_T4_His | + | ++ |
| Spike_ΔCter_Linkers_T4_His | - | + |
| Spike_ΔCter_ΔFurin_Linkers_T4_His | + | ++ |
| Spike_ΔCter_2P_T4_His | - | + |
| Spike_ΔCter_ΔFurin_2P_T4_His | + | +++ |
| Spike_ΔCter_2P_Linkers_T4_His | - | + |
| Spike_ΔCter_ΔFurin_2P_Linkers_T4_His | + | ++ |
| Spike_RBD_His | +++ | n.a. |

### 3.2. CHO expression and purification of SARS-CoV-2 proteins

Stable recombinant cell pools expressing S trimer and RBD were generated using puromycin selection. From a total of 300 clonally derived cell populations, those with the highest expression levels of S trimer and RBD were expanded and a fed-batch production process was developed. Cell culture process conditions, such as medium formulations, feeds, feed and temperature shift timing were evaluated at 10mL scale in TPP^®^ TubeSpin 50 bioreactor tubes as previously described [13, 14]. Optimal conditions were selected on the basis of product yield, product quality, viable cell density (VCD) and cell viability. The production process was scaled up to 200 mL shake flasks, to 10 L and to 40 L stirred tank bioreactors (STR). Viable cell density (VCD) and viability remained high for at least 10 days (Figure 2). In addition, viable cell density (VCD) and viability profiles were comparable between shake flasks, 10L and 40L STRs, indicating process scalability. From ExcellGene’s experience in developing high-yielding manufacturing processes for clinically relevant recombinant antibodies and other proteins since almost 20 years, there is high confidence to assure feasibility to any scale of operation, up and beyond the 2000 Liter scale. Krammer [15] rightly points to expression and scale-up difficulties for production large and complex proteins in global SARS-CoV-2 vaccines, such as the S trimer. However, since the mid 1980s CHO cells have delivered hundreds of kilograms and tons of proteins from bioreactors at scales of up to 20’000 L [16]. One leading subunit vaccine candidate similar to the one presented here includes the transmembrane region of the Spike trimeric protein and is produced and purified from Baculovirus infected insect cells (SF-9). To the knowledge of the authors, this technology has not been scaled up to any scale larger than 100 L. This vaccine may face a serious manufacturability issue, when intended for global use. Also, insect cells present a different glycosylation pattern on proteins (pauci-manosidic, alpha 1-3 fucose) than mammalian cell systems [17]. One could argue that a manufacturing system with more human like glycosylation patterns could be a superior vaccine, since it mimics more closely the authentic protein. This is particularly important since each Spike monomer has 9 predicted N- and 3 O-glycosylation sites [18] that reveal the polypeptide backbone for potential antigenic epitopes only at a part of its surface [19].

**Figure 2.**
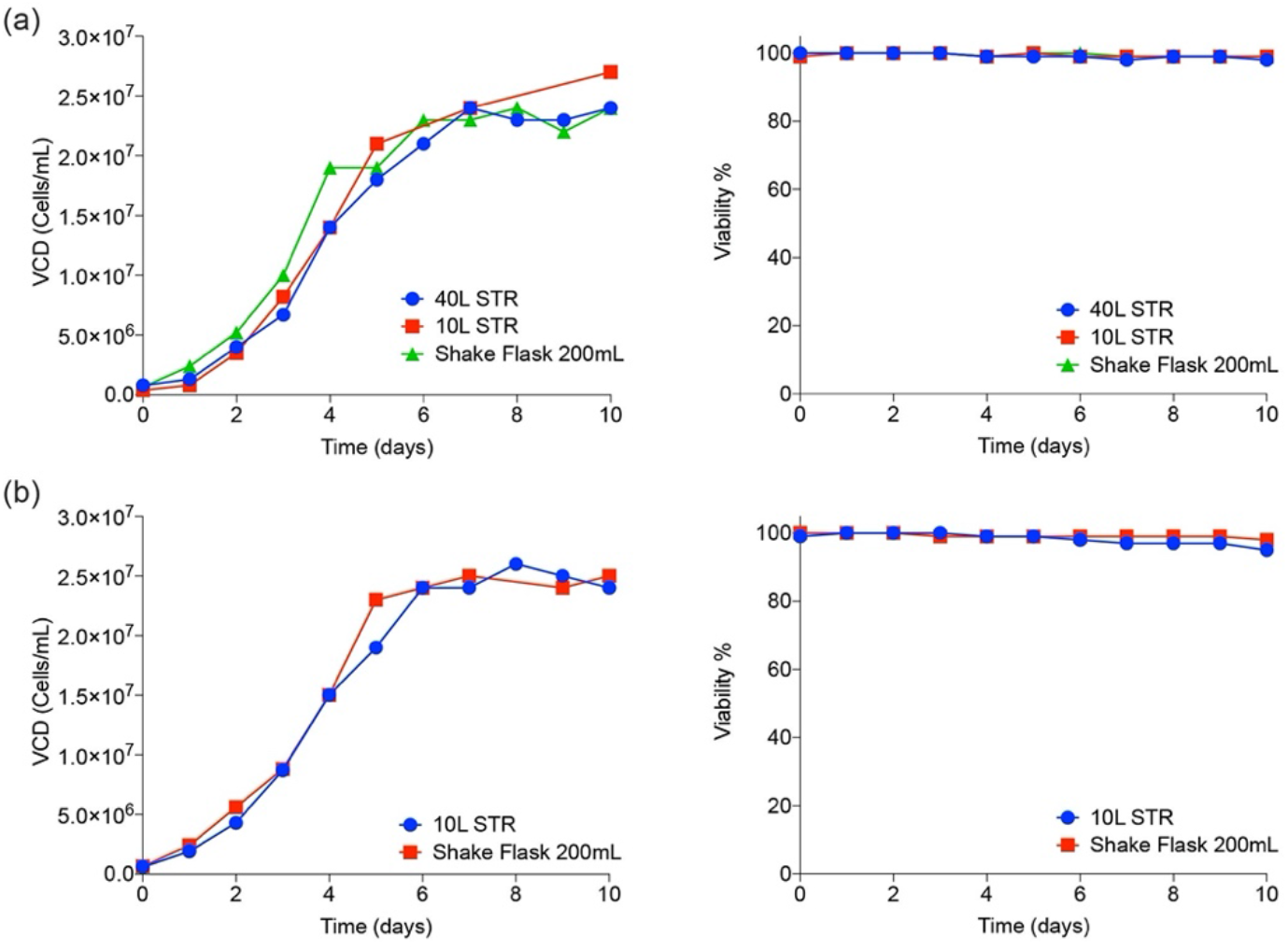
Cell culture scale-up performance in shake flasks and Bioreactors. Viable cell densities (cells/mL) and cell viability (%) are shown at various scales of operation for both S trimer (a) and RBD (b).

### 3.3. Molecular characterization and electron microscopy analysis of Trimeric Spikes

S trimer and RBD from cell-free culture supernatant were purified using an immobilized metal affinity chromatography (IMAC) capture step followed by preparative size exclusion. Eluates were concentrated and formulated at 1mg/mL in PBS pH7.4. Purity of the final products was estimated to be over 95% as analyzed by HPLC-SEC (Figure 3 a and b).

**Figure 3.**
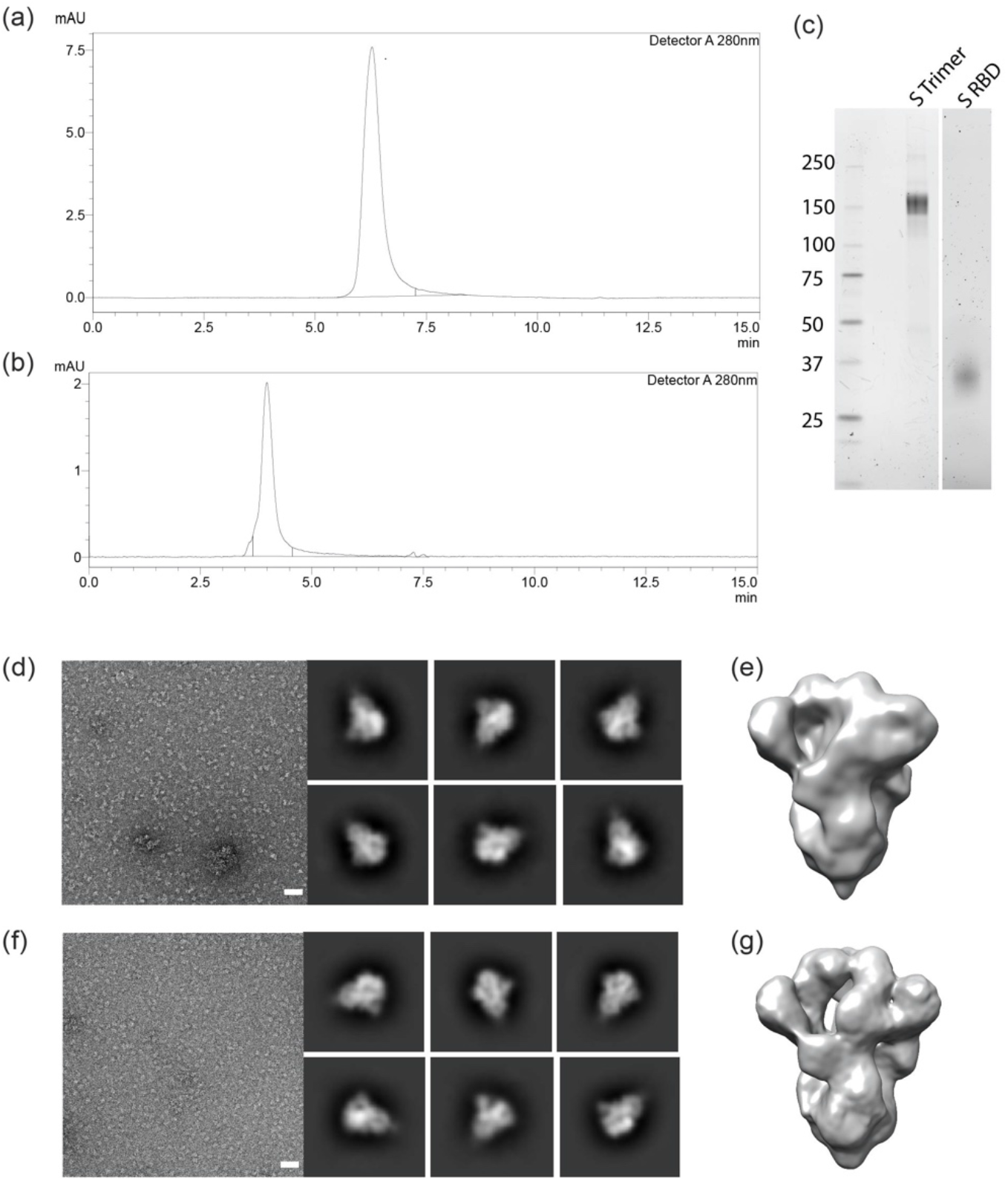
Characterization of the recombinant trimeric S trimer and RBD. Size-exclusion chromatography (SEC) analysis plot with a single peak at 4.0 minutes and 6.4 minutes. SDS-PAGE gel showing bands for S trimer at different dilutions (c) and (d) for RBD monomer, NR=non-reduced, R=reduced. Negative stain electron microscopy of SARS-CoV-2 S trimer (Wuhan, d,e, D614G, f-g). White bars in d and f: 50 nm. Six 2D class averages are shown to the right of each representative micrograph. 3D reconstructions are shown in (e) and (g).

HPLC-SEC (non-denaturing conditions) indicates that the S protein complex is trimeric (about 460 kDa) and the RBD is monomeric. The expected size of 150-160 kDa and 29 kDa for the Spike and RBD monomers, respectively, were confirmed by reducing SDS-PAGE (Figure 3 c, d). The heavy glycosylation of S-trimer contributes significantly to size heterogeneity, as has been seen with other CHO cell produced virus derived proteins, such as truncated surface proteins from Ebola and HIV. Presently ongoing work will characterize the glycosylation of the S-trimer from the CHO cell process. Trimeric confirmation was also confirmed using negative-stain electron microscopy (Figure 3 e - h).

### 3.4. Inhibition of SARS-CoV-2 infection by SARS-CoV-2-proteins

SARS-CoV-2 virus was mixed with serially diluted RBD or Spike trimer and the mixture was inoculated on Vero E6 cells. 48 hours later, SARS-CoV-2 infection was visualized and quantified (Figure 4 a and b). The presence of both the RBD Spike fragment and Spike trimer reduced infection of Vero cells. For RBD, reduction of infectivity was observed at a concentration of 20 *μ*M and higher. For the S trimer (both Wuhan and D614G), a near 100-fold lower concentration reduced infection (Figure 4 b). Taken together, these results show that CHO-produced trimeric S and RBD are able to compete with SARS-CoV-2 for binding to host cells. No differences were observed between the 614D (Wuhan) and the D614G variants of the Spike trimer.

**Figure 4.**
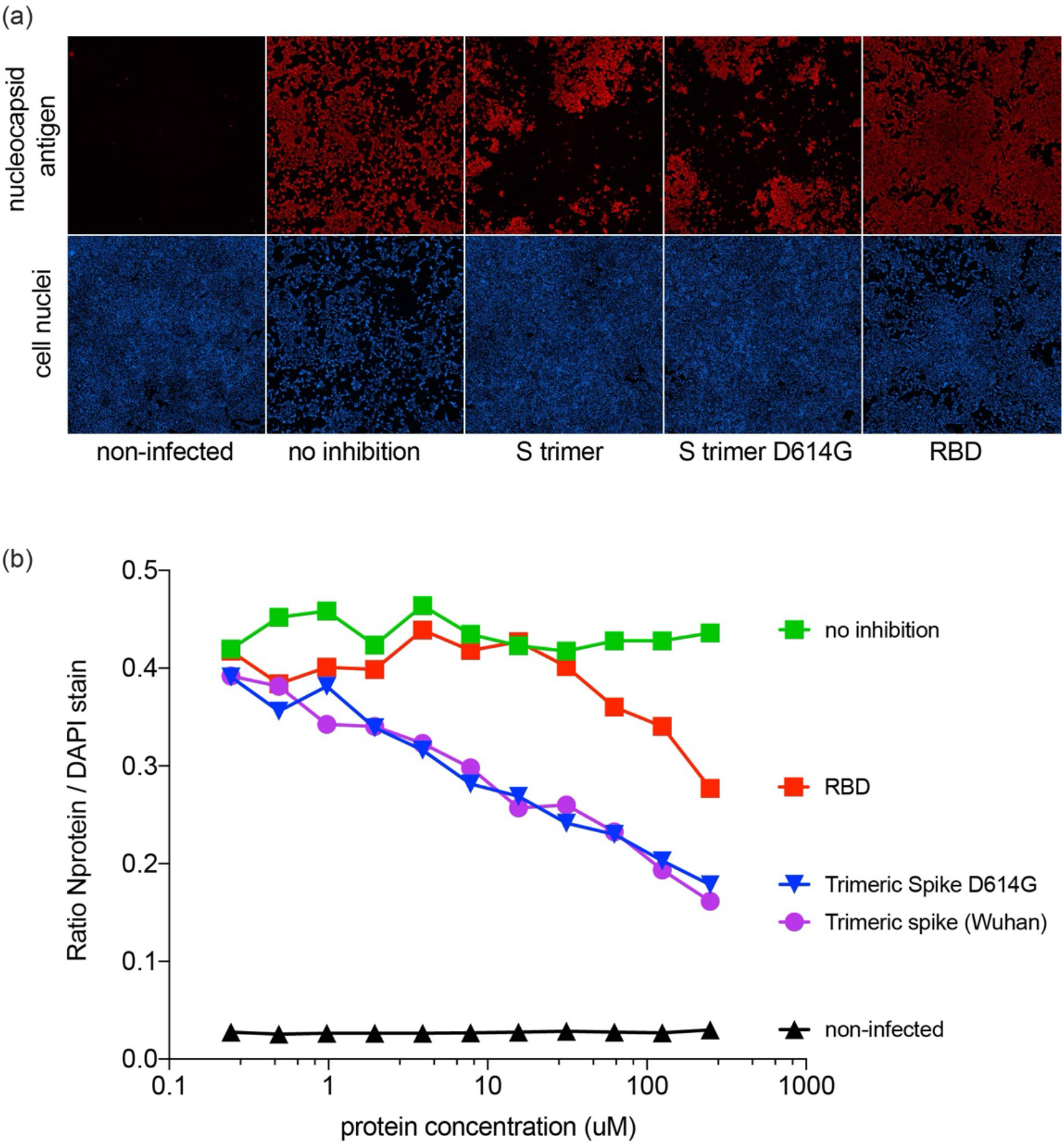
Inhibition of SARS-CoV-2 infection in Vero E6 cells (a) Inhibitory effect of S trimer and RBD on infection of Vero E6 cells. SARS-CoV-2 antigen-positive cells were visualized by immunofluorescent staining. Virus nucleocapsid antigen staining (red), cell nucleus staining (blue). All methods as described previously[10]. (b) Inhibitory effect of S trimer and RBD on infection of Vero E6 cells. SARS-CoV-2 antigen-positivity fluorescence was quantified as described [10, 11].

### 3.5. Sensitivity and specificity of SARS CoV-2 Spike proteins for detection of antibodies

The trimeric Spike and RBD proteins were used to detect SARS-CoV-2-specific IgG in a semi-quantitative bead-based serological assay [10]. We specifically selected a set of COVID-19 serum samples that cover a large range of SARS-CoV-2-specific antibody concentrations. Sera (n = 72) were collected from PCR-confirmed nonhospitalized (n = 63) and hospitalized (n = 9) COVID-19 patients between days 4 and 40 after disease onset. Control sera (n = 79) were collected before the emergence of SARS-CoV-2 from patients with seasonal corona-induced Influenza-like illness (n = 44), non-corona, Influenza-like illness (n = 29), and from healthy individuals (n = 6).

For all proteins, the median signal intensity for the COVID samples was over 400-fold higher than for the control samples (Figure 5 a). Receiver Operator Characteristic (ROC) analysis (Figure 5 b) showed that trimeric Spike (both Wuhan and D614G) provided higher test-sensitivity and higher specificity characteristics compared to both S1 monomer and commercial RBD produced in HEK293 (p = 0.03; 0.04), and RBD produced in CHO (p = 0.05). Statistical analysis [20, 21] showed no differences between the ROC of S1 monomer and both RBD’s, neither compared to Wuhan nor to D614G trimeric Spikes. Taken together, our results indicate that CHO-produced Trimeric Spike can deliver higher diagnostic specificity and sensitivity than S1 monomer and RBD. These observations are in-line with previous reports [3].

**Figure 5.**
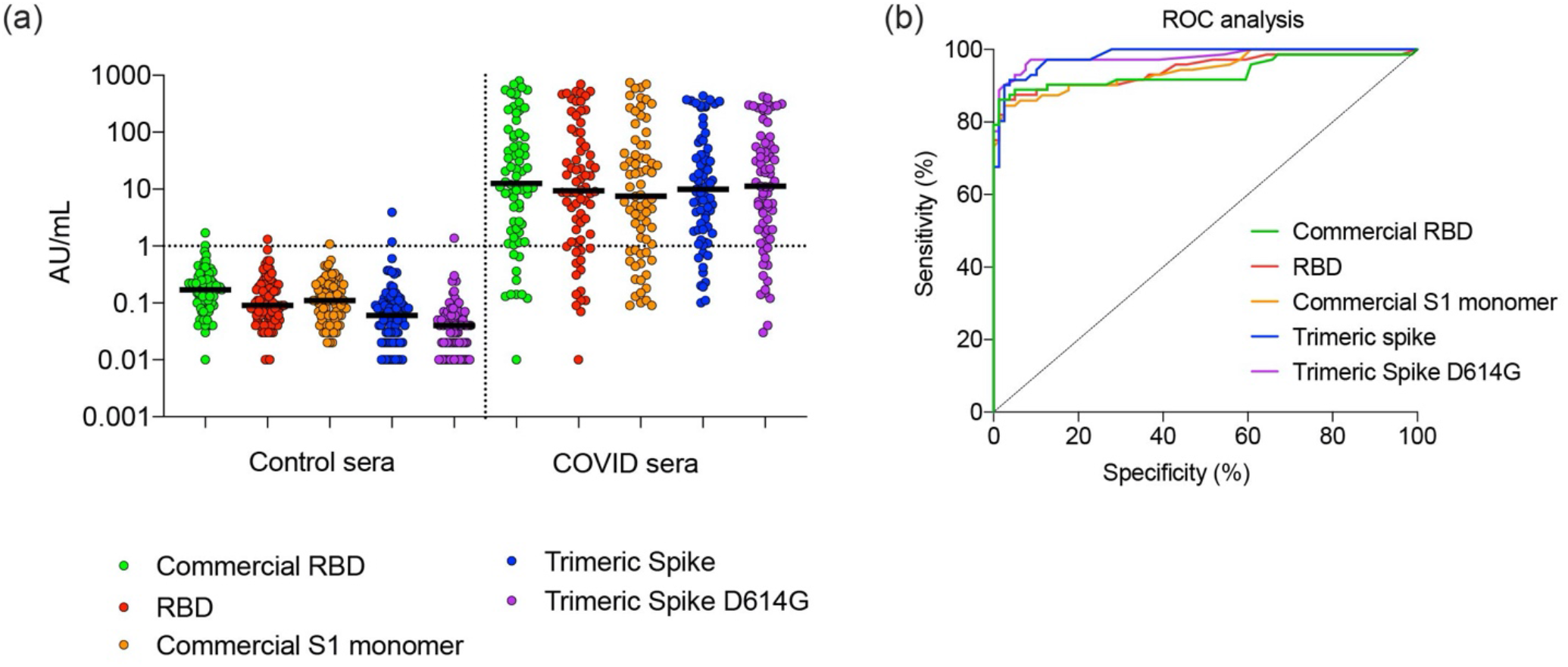
Performance of RBD, monomeric S1 and S trimer in a diagnostic assay to identify COVID-19 patients. (a) Control sera (n = 151) and COVID-19 sera (n = 72) collected at day 4-40 of symptoms were tested and compared for concentrations of IgG. Median concentration and 95% confidence intervals are shown. (b) The sera tested in (a) were analyzed by ROC. Abbreviations: AU, arbitrary unit; RBD, receptor binding domain; ROC, receiver operator characteristic; S1, Spike protein subunit 1.

## 4. Conclusion

Using an optimized CHO expression system, and a scalable, chemically defined production process, trimeric SARS CoV-2 Spike proteins with mammalian-type glycosylation could be provided in sufficient quantities. Analysis of the resulting protein shows that it is of high purity and trimeric. Functional analysis showed it to be efficient in blocking virus infectivity in an in-vitro model. In addition diagnostic performance of the trimeric Spike for SARS-CoV-2 specific IgG was shown to exceed that of RDB and monomeric S1 protein.

## Author Contributions

Conceptualization, P.P., D.K., M.J.W., J.K. and F.M.W.; methodology, P.P., J.K., R.D., V.A., G.P., J.S.M., D.K., M.J.W.; validation, P.P., J.K., D.K., M.J.W.; molecular analysis and EM work, D.K., J.S.M, P.O.B.; reactivity toward sera, P.R., R.D., G.dH.; formal analysis, F.M.W., P.P., J.K., J.S.M, R.D.; writing—first draft preparation, F.M.W.; writing—review and editing, P.P., J.K., F.M.W.; supervision, P.P., J.K., D.K.; project administration, M.J.W., D.K.. All authors have read and agreed to the published version of the manuscript.

## Acknowledgements

Frederico Pratesi and Paola Migliorini (Clinical Immunology Laboratory, University of Pisa, Italy, are thanked for rapid and early provisioning of Covid-19 sera providing encouraging data to the team towards further and more profound work on this program.

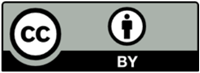 © 2020 by the authors. Submitted for possible open access publication under the terms and conditions of the Creative Commons Attribution (CC BY) license (http://creativecommons.org/licenses/by/4.0/).

